# pyngoST: fast, simultaneous and accurate multiple sequence typing of *Neisseria gonorrhoeae* genome collections

**DOI:** 10.1101/2023.10.23.563537

**Authors:** Leonor Sánchez-Busó, Andrea Sánchez-Serrano, Daniel Golparian, Magnus Unemo

## Abstract

Extensive gonococcal surveillance has been performed using molecular typing at global, regional, national and local levels. The three main genotyping schemes for this pathogen, Multi-Locus Sequence Typing (MLST), *Neisseria gonorrhoeae* Multi-Antigen Sequence Typing (NG-MAST) and *N. gonorrhoeae* Sequence Typing for Antimicrobial Resistance (NG-STAR), allow inter-laboratory and inter-study comparability and reproducibility and provide an approximation to the gonococcal population structure. With high-throughput whole-genome sequencing (WGS), we obtain a substantially higher and more accurate discrimination between strains, i.e., compared to previous molecular typing schemes where isolates with the same sequence type often have a different genomic background. However, WGS remains unavailable or not affordable in many laboratories, accordingly, bioinformatic tools that allow the integration of data among laboratories with and without access to WGS is imperative for a joint effort to increase our understanding of global pathogen threats.

Here, we present pyngoST, a command-line Python tool for a fast, simultaneous and accurate sequence typing of the WHO priority pathogen *N. gonorrhoeae*, from WGS assemblies. pyngoST integrates MLST, NG-MAST and NG-STAR, and can also designate NG-STAR clonal complexes and NG-MAST genogroups, facilitating multiple sequence typing from large WGS assembly collections. Exact matches for existing alleles and STs are reported, but also closest matches of new alleles and STs. The implementation of a fast multi-pattern searching algorithm allows pyngoST to be rapid and report results on 500 WGS assemblies in under 1 minute. The mapping of typing results on a core genome tree of 2,375 gonococcal genomes revealed that NG-STAR is the scheme that best represents the population structure of this pathogen, emphasizing the role of antimicrobial use and antimicrobial resistance (AMR) as a driver of gonococcal evolution.

**IMPACT STATEMENT:** Molecular typing has been key for *N. gonorrhoeae* epidemiological and AMR surveillance, and WGS has revolutionized this typing. The most frequently used molecular typing schemes include MLST, NG-MAST and NG-STAR, and modifications of those. These schemes can be extracted from WGS assemblies for comparability and reproducibility of results with laboratories that do not have access to WGS technologies. pyngoST is a unique command-line Python tool that integrates all these common typing schemes under the same framework and performs rapid simultaneous user-defined multiple typing of large number of gonococcal genomes through a fast multi-pattern searching algorithm. Typing results on 2,375 gonococcal genomes revealed that NG-STAR best represents the genomic population structure of *N. gonorrhoeae*, highlighting the importance of antimicrobial use and AMR on the evolution of this pathogen.

**DATA SUMMARY:**

1. pyngoST is written in Python 3 and is available from Github under the GPL-3.0 License (https://github.com/leosanbu/pyngoST).
2. The script can be installed via the Python ‘pip’ package.
3. Genome assemblies used in this study are from the Euro-GASP 2018 WGS survey and are available from Pathogenwatch: https://pathogen.watch/collection/eurogasp2018 (1,2). Pairwise single nucleotide polymorphism (SNP) distances among isolates from this dataset are also available from the same link.
4. Results from running pyngoST on the Euro-GASP 2018 WGS dataset can be explored from Microreact: https://microreact.org/project/wYpBzCs9A6Uf7HEMA6zmmY-eurogasp2018-pyngost.

## INTRODUCTION

Molecular typing of bacterial species revolutionized epidemiological surveillance of pathogens in the late 20th century, especially after the development of DNA-based techniques such as Polymerase Chain Reaction (PCR) amplification. In the 1990s, Multi-Locus Sequence Typing (MLST) (3) emerged as a highly standardized molecular typing method that involved the PCR amplification and traditional Sanger DNA sequencing of fragments of several housekeeping genes to define unique allele profiles for bacterial strains. The different allelic profiles are subsequently designated as divergent sequence types (STs). This methodology greatly enhanced our understanding of bacterial pathogen epidemiology, antimicrobial resistance (AMR) and surveillance (4).

For the sexually-transmitted human pathogen *Neisseria gonorrhoeae*, three different sequence-based typing schemes have been used most frequently (Table 1): *Neisseria* spp. MLST (5), *N. gonorrhoeae* Multi-Antigen Sequence Typing (NG-MAST) (6), and *N. gonorrhoeae* Sequence Typing for Antimicrobial Resistance (NG-STAR) (7). The *Neisseria* spp. MLST scheme was developed to characterize molecular lineages of *N. gonorrhoeae, N. meningitidis* and *N. lactamica* and examines fragments of seven slowly evolving housekeeping genes (*abcZ, adk, aroE, fumC, gdh, pdhC* and *pgm*) (5). In contrast, NG-MAST examines fragments of two highly variable genes, *porB* and *tbpB*, encoding the PorB (Porin B) and TbpB (Transferrin-binding protein B) outer membrane proteins and it was developed to enhance strain discrimination (6). Due to its limitation of examining only two highly-variable loci, often including recombination events, NG-MAST is typically not representative of the population structure of *N. gonorrhoeae*. However, NG-MAST STs can be grouped into genogroups following a set of similarity rules between the alleles of different isolates to improve the representation of this population structure (8,9). However, NG-MAST genogroups are dataset-specific, ST-genogroup relations are not stable over time and thus, genogroups are not comparable between studies or time points (1,8). Alternative methods are recommended, such as NG-STAR, for a more robust and transferable strain characterization that also reflects AMR.

**Table 1.** Molecular typing schemes implemented in pyngoST.

| Scheme (source) | Number of loci | Number of STs* | Loci (number of alleles)* | Sequence lengths (bases) |
| --- | --- | --- | --- | --- |
| NG-STAR (7)<br>(ngstar.canada.ca) | 7 | 5,290 | *penA* (n=615)<br>*mtrR* (n=593)<br>*porB* (n=73)<br>*ponA* (n=20)<br>*gyrA* (n=62)<br>*parC* (n=187)<br>*23S rDNA* (n=82) | 1,745-1,752<br>539-719<br>30<br>74-76<br>263-264<br>331-334<br>566-571 |
| MLST (5,22)<br>(PubMLST) | 7 | 17,626 | *abcZ* (n=1,257)<br>*adk* (n=1,003)<br>*aroE* (n=1,280)<br>*fumC* (n=1,354)<br>*gdh* (n=1,331)<br>*pdhC* (n=1,254)<br>*pgm* (n=1,341) | 432-435<br>464-467<br>481-493<br>463-467<br>500-513<br>479-503<br>447-489 |
| NG-MAST v2<br>(6,22)**<br>(PubMLST) | 2 | 22,047 | *porB* (n=13,051)<br>*tbpB* (n=3,183) | 394-526<br>367-437 |
| **Total (profiles and alleles)** |  | **44,963 STs** | **26,686 alleles**<br>**(53,372 forward and reverse)** |  |

\*Number of STs and alleles of each locus according to a database downloaded in August 2023 (11/08/2023).
\*\*Note that the optimised scheme NG-MAST v2 from PubMLST is used by default.
NG-STAR: *N. gonorrhoeae* Sequence Typing for Antimicrobial Resistance. MLST: Multi-Locus Sequence Typing. NG-MAST: *N. gonorrhoeae* Multi-Antigen Sequence Typing

Monitoring AMR gonococcal lineages has been stated as crucial to combat AMR gonococci (1,8,10). The NG-STAR scheme was developed as a standardized AMR typing tool that combined strain typing with AMR profiling (7). NG-STAR examines sequence fragments of seven AMR determinants associated with resistance or decreased susceptibility to antibiotics, such as penicillin, ciprofloxacin, the Extended-Spectrum Cephalosporins (ESCs) cefixime or ceftriaxone, and azithromycin: *penA, mtrR, porB, ponA, gyrA, parC* and 23S rRNA. This typing scheme allows monitoring the emergence and spread of lineages with different degrees of antimicrobial susceptibility or AMR. NG-STAR STs can be grouped into Clonal Complexes (CC) for a more consistent fit with the *N. gonorrhoeae* population structure, simplifying nomenclature and providing reproducibility into AMR lineage definition (11).

Despite MLST being a breakthrough for pathogen epidemiology, the real revolution arrived with whole genome sequencing (WGS) (12,13), which provided a comprehensive view of genomes, enabling high-resolution strain typing, phylogenetic analysis, and detailed characterization of genetic variations and AMR determinants. The progressive decrease in the cost of high throughput short-read sequencing has made WGS a cost-effective tool compared to performing multiple PCR reactions followed by Sanger sequencing for strain typing. However, molecular typing tools are still in use in public health laboratories because they are used for simplified naming of strains/clones/clades and they are included in standardized protocols to ensure inter-laboratory reproducibility and quality of the results. The standardization of WGS-based protocols for epidemiological surveillance of pathogens is still undergoing, as it requires specialized infrastructure and personnel training (14).

Traditionally, allele sequences and allelic profiles are manually submitted to web servers, revised and assigned by data curators. With the appropriate bioinformatic tools, molecular typing alleles can be extracted from bacterial genomes. To facilitate molecular typing from collections of genome sequences, command-line tools have been developed, such as mlst (15), SRST2 (16), ARIBA (17), or NGMASTER, which is specific for the NG-MAST scheme (18). The rules to calculate NG-MAST genogroups have been reported in previous studies (8,9), however, there is still no publicly-available tool that implements them. Finally, command-line tools that automatically perform large-scale NG-STAR or NG-STAR CCs typing are lacking.

Here, we present pyngoST, a python tool that unifies molecular typing of *N. gonorrhoeae* from genome assemblies. pyngoST implements the Aho-Corasick (AC) multi-string searching algorithm for an ultrafast simultaneous screening of all loci included in the MLST, NG-MAST and NG-STAR schemes on the input assembly files. pyngoST can also output NG-STAR CCs and calculate NG-MAST genogroups. We also provide an evaluation of the concordance of these typing schemes with the gonococcal population structure obtained from WGS.

## METHODS

### Implementation of pyngoST

pyngoST has been modularly implemented in Python 3.8 under a GNU GPLv3 license and is available from Github (https://github.com/leosanbu/pyngoST). Installation can be performed from the source code or from Pypi using *pip*. pyngoST is written as an open-source tool containing a main script *pyngoST*.*py*, where the principal functions are contained in and called from *pyngoST_utils*.*py*. The main dependencies include the *pyahocorasick* module (https://github.com/WojciechMula/pyahocorasick) and Biopython (19). *pyahocorasick* is a fast and memory efficient library written in C that contains the *ahocorasick* Python module for multi-pattern string search via the AC algorithm.

The usage of pyngoST requires the building of a local database, from which three different modules can be run (Figure 1): 1) molecular typing of *N. gonorrhoeae* from assemblies in fasta format by any of MLST, NG-MAST, NG-MAST genogroups, NG-STAR and NG-STAR CCs, 2) sequence typing from multiple allelic profiles of any of MLST, NG-MAST, NG-STAR and NG-STAR CCs contained on a tabular or comma-separated table, and 3) classification of NG-STAR STs into NG-STAR CCs from a tabular or comma-separated table.

**Figure 1.**
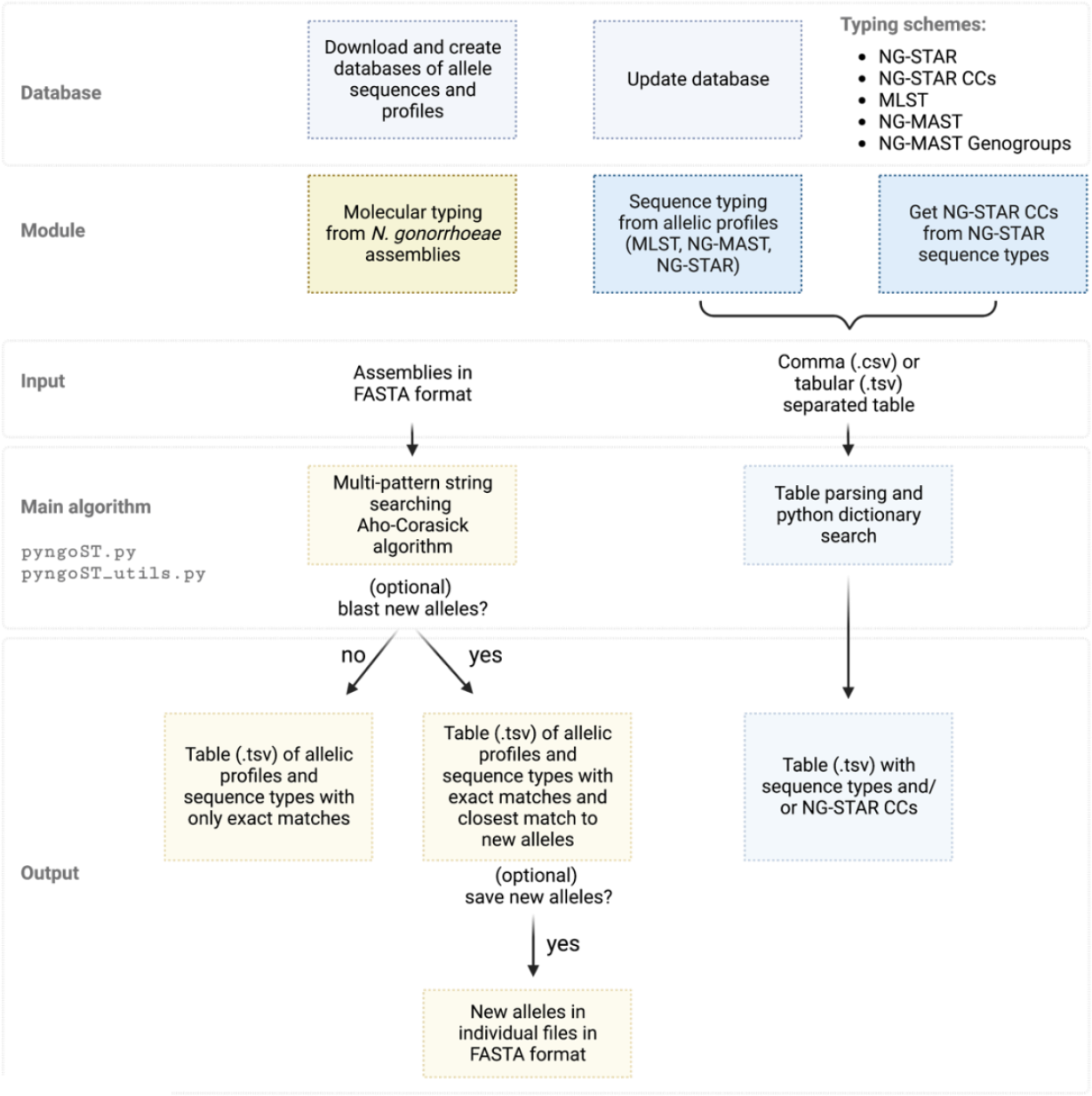
Structure of the pyngoST algorithm. From a database of allele sequences and sequence type (ST) profiles, pyngoST performs simultaneous molecular typing of *Neisseria gonorrhoeae* assemblies using the multi-pattern string searching Aho-Corasick algorithm through three schemes: *N. gonorrhoeae* Sequence Typing for Antimicrobial Resistance (NG-STAR) (7), Multi-Locus Sequence Typing (MLST) (5) and *N. gonorrhoeae* Multi-Antigen Sequence Typing (NG-MAST) (6) (Table 1). NG-STAR Clonal Complexes (CCs) (11) and NG-MAST genogroups (8,9) can also be reported. Closest matches of new alleles to existing sequences can be computed using BLAST (20) and these can be saved for manual submission by the user to the corresponding web servers. pyngoST can also be used to only obtain NG-STAR CCs from NG-STAR STs according to the database as well as to report STs from a table of allelic profiles of any of the MLST, NG-MAST and NG-STAR schemes.

NG-MAST genogroups are calculated following previously-described rules: NG-MAST profiles are assigned into a genogroup if one identical allele is shared and the other allele shows ≥99% similarity (≤5 bp difference for *porB* and ≤4bp difference for *tbpB*) or the concatenated sequence of both alleles shows ≥99.4% similarity to the concatenated sequence of both alleles of the main ST in the genogroup (1,8,9).

Novel allele sequences of any of the three main schemes (MLST, NG-MAST and NG-STAR) can be optionally assigned to the closest match in the database by using a local installation of BLASTn (20). These new sequences can be extracted from the assemblies and saved as fasta files so the user can submit them to the corresponding web server for assignment by a data curator. Large data collections can benefit from multithreading this task through the implementation of a ThreadPoolExecutor via the *concurrent*.*futures* Python module (21).

### Database building

The usage of pyngoST relies on the building of a local database, which can be downloaded or updated using the -d (calls the *download_db()* function) and -u (calls the *update_db()* function) options, respectively. MLST (PubMLST scheme ID = 1, 7 loci) and NG-MAST (PubMLST scheme ID = 71, NG-MAST v2, 2 loci) allele sequences and ST profiles are downloaded from PubMLST https://pubmlst.org, while NG-STAR allele sequences of the 7 loci and the assigned ST profiles are downloaded from https://ngstar.canada.ca (Table 1). Allele sequences and ST profiles are stored in python dictionaries. These are used to create an AC automaton (*make_ACautomaton()* function) using the *pyahocorasick* module, which is encapsulated into a *pickle* file that is loaded in subsequent pyngoST runs to ensure speed in database loading. As reference, a database downloaded in August 2023 contained 44,963 STs and 26,686 alleles (Table 1). As draft assemblies are often used, the reverse complementary sequence of each allele is obtained and stored with the database, which contained 53,372 sequences.

Note that the web server of the original NG-MAST scheme (6) is no longer available and that a refined and optimised version, NG-MAST v2, curated and maintained in PubMLST (22), is used by default by pyngoST. This improved version performs an additional curation of *porB* and *tbpB* sequences to account for the hyper variability found in this scheme. However, older versions of the latest available NG-MAST database could be manually included by the user instead of this optimised version. This is important as it can lead to some inconsistencies when comparing NG-MAST v2 results with the original NG-MAST results from previous studies.

NG-STAR CCs are obtained by applying a full goeBurst algorithm on an up-to-date NG-STAR database and grouping those as previously described (11). NG-STAR CCs are integrated into the database at the time of building with the -cc option (calls the *integrate_ngstar_ccs*() function) by providing a comma-separated file containing NG-STAR STs and the associated CCs. There is still not a regular publicly-available update of NG-STAR CCs, but the latest run (from May 2023) is available from the pyngoST Github repository, and will be posteriorly further updated and shared with the scientific community. If the database files are manually modified (i.e. by adding new STs or allele sequences), the AC automaton and *pickle* file can be updated with -u. If NG-STAR CCs were not provided at the time of the database building, they can be provided when updating.

### Tests of performance

Ten random genome sets of different sizes (25, 50, 100, 200, 400, 800 and 1,600 genomes) were obtained from the Euro-GASP 2018 dataset (n=2,375 genomes) (1). The performance of pyngoST was evaluated by computing the run time of different combination of parameters: i) only retrieving exact allele matches from the database for MLST, NG-MAST, NG-STAR and NG-STAR CCs, ii) blasting new alleles for MLST, NG-MAST and NG-STAR and reporting the closest alleles, iii) computing NG-MAST genogroups from different data set sizes and iv) using multithreading. All tests were run on a MacBook Pro 2,3 GHz Intel Core i7 laptop with 4 cores, 8 threads and 16 GB RAM.

### Representation of the gonococcal population structure

Genome assemblies and a core genome MLST (cgMLST) phylogenetic tree from 2,375 isolates from the European Gonococcal Antimicrobial Surveillance Programme (Euro-GASP) 2018 WGS study were downloaded from Pathogenwatch (https://pathogen.watch/collection/eurogasp2018) (1,2). pyngoST was run on the assemblies to obtain MLST STs, NG-MAST STs, NG-MAST genogroups, NG-STAR STs and NG-STAR CCs. A consistency and retention index (CI and RI) were calculated using the *phangorn* v2.7.1 R package to estimate the fitness of each scheme to the phylogenetic tree under a parsimony model. The CI measures the relative homoplasy of the data and the RI measures monophyly. Values of 1 represent a perfect fit. Pairwise single nucleotide polymorphisms (SNPs) in the core genome among the isolates of the Euro-GASP 2018 WGS dataset were also obtained from Pathogenwatch (1,2). and processed in R to evaluate the distance between isolates belonging to the same sequence types, CCs and genogroups.

## RESULTS AND DISCUSSION

*N. gonorrhoeae* is one of the top WHO priority pathogens due to AMR (23,24) and the results of epidemiological surveillance that use either molecular typing or genomic data are constantly being published, justifying the need of tools that rapidly and accurately integrate both types of data. Here, we present pyngoST (Figure 1), a command-line based python tool that unifies molecular typing for *N. gonorrhoeae*, for which several typing schemes have been used to discriminate among lineages since before the genomic era. Initially, typing relied on time-consuming amplification-based protocols followed by capillary sequencing of a panel of housekeeping (MLST) or highly-variable genes (NG-MAST). More recently, sequencing of a panel of genes associated with AMR (NG-STAR) was shown to represent main genomic lineages similarly to MLST but with the benefit of additionally representing AMR profiles. Modifications of NG-STAR and NG-MAST have also been used in the form of NG-STAR CCs and NG-MAST genogroups, respectively. With the release of pyngoST, we provide a framework for the unification of these five typing schemes that can be used in a fast, memory-efficient and accurate manner on large collections of genome assemblies. pyngoST can also define STs and identify NG-STAR CCs from tables containing allelic profiles, being also a useful tool for integrating typing of *N. gonorrhoeae* in laboratories without access to WGS and high-performance computing.

### Fast, simultaneous and accurate molecular typing from genome assemblies

pyngoST was conceived to be fast when run even in large collections of genome assemblies. It uses a simultaneous multi-string searching algorithm for exact matching of a database of allelic sequences in direct and reverse orientation. The database used at the time of preparing this manuscript contained 53,372 sequences representing the forward and reverse sequences of all the alleles in the 7 MLST, 2 NG-MAST and 7 NG-STAR loci. We ran pyngoST on the 2,375 genomes of the Euro-GASP 2018 genomic survey (1) on a laptop computer and it took 3.84 minutes to accurately report the profiles and STs of the three schemes plus the NG-STAR CCs using exact matches to existing alleles and STs in the database. Using the -b option to obtain the closest known alleles to new alleles using BLASTn on the whole dataset pyngoST took 11.87 minutes. However, in this mode, the run time will be dependent on the number of new alleles in the dataset. Ten random genome sets of different sizes (25, 50, 100, 200, 400, 800 and 1,600 genomes) from the same study (1) were used to evaluate the performance of pyngoST in terms of speed and a linear increase in run time with an increasing number of genomes was observed. Computation time ranged from a minimum of 3.2 seconds for a 25-genome dataset to 2.3 minutes for a 1,600-genome dataset. Requesting the closest known alleles to new sequences (-b) resulted in minimum a computation time of 15.7 seconds in a dataset of 25 genomes and a maximum computation time of 18.8 minutes in a dataset of 1,600 genomes. If different isolates have identical alleles that have not been identified before, they are assigned the same number. Obtaining the closest known alleles is a more time-consuming task than only reporting exact alleles and can be parallelized using multithreading, which reduces runtime proportionally to the number of threads requested. To further streamline the submission process for the user, these new sequences can be exported to a fasta file using the -a option, making it easier to submit to the hosting web server for assignment. NG-MAST genogroups can also be requested from the NG-MAST results (-g option). This task was performed after typing and took 31.8 minutes for the complete dataset of 2,375 genomes. For different dataset sizes, run time increased linearly with size, with an average of 4.1 minutes for 25 genomes to an average of 16.5 minutes for 1,600 genomes.

### NG-STAR best represents gonococcal population structure

Previous comparisons on the distribution of gonococcal STs or CCs obtained from molecular typing on gonococcal phylogenomic trees have shown decent concordance values (8,11). In this study, using a collection of 2,375 genomes from the Euro-GASP 2018 genomic survey, we observed that the scheme that best fitted the phylogenomic tree topology (Figure 2a) was NG-STAR (RI=0.949, CI=0.819), followed by MLST (RI=0.962, CI=0.683), NG-STAR CCs (RI=0.945, CI=0.476), NG-MAST (RI=0.793, CI=0.607) and NG-MAST genogroups (RI=0.827, CI=0.413). Singleton STs, however, inflate these estimates, as they can fit on any topology, and this dataset contains 39% of singletons for MLST (64 out of 164) and over 50% of singletons for NG-STAR (214 out of 418 STs) and NG-MAST (245 out of 483). Better estimates have been previously described for the different schemes (8,11), however, several factors may affect these calculations, such as the composition of the dataset or the accuracy of the phylogenomic tree. Low CIs were obtained from NG-STAR CCs and NG-MAST genogroups, indicating high homoplasy of these classification schemes in the tree. This can be expected as they group different NG-STAR STs and NG-MAST STs, respectively, following specific rules on the target loci. Genomic variability in the rest of the core genome due to mutation or recombination often splits isolates of the same ST in different genomic lineages on the phylogenetic tree. This fact is probably aggravated for NG-STAR CCs and NG-MAST genogroups. Nonetheless, the RI, which estimates monophyly of the classification on the tree, was very high for NG-STAR CCs, NG-STAR STs and MLST and this is reflected on the mapping over the phylogenomic tree (Figure 2a).

**Figure 2.**
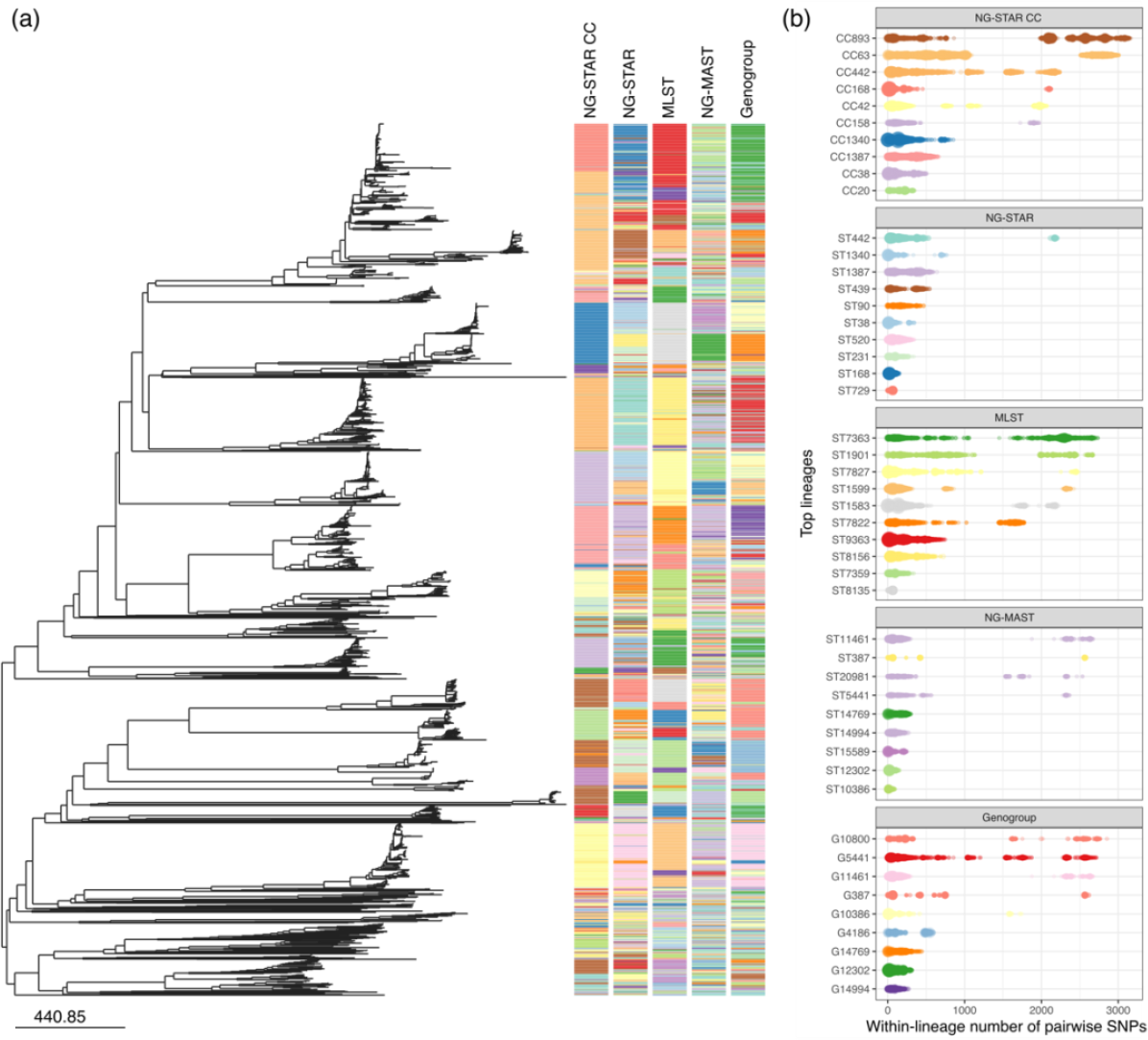
Representation of the *Neisseria gonorrhoeae* genomic population structure through the existing molecular typing schemes. (a) Phylogenetic tree of the Euro-GASP 2018 genomic dataset (1) as calculated from Pathogenwatch from SNPs in the core genome (2) and represented in Microreact (25). Coloured strips next to the tree represent, in the following order, different *N. gonorrhoeae* Sequence Typing for Antimicrobial Resistance Clonal Complexes (NG-STAR CCs) (11), NG-STAR Sequence Types (STs) (7), Multi-Locus Sequence Typing (MLST) STs (5,22), *N. gonorrhoeae* Multi-Antigen Sequence Typing (NG-MAST) STs (6,22) and NG-MAST genogroups (8,9), as obtained through pyngoST. Colours were automatically assigned by Microreact. (b) Number of pairwise single nucleotide polymorphisms (SNPs) in the core genome of the top ten lineages of each of the schemes: NG-STAR CCs, NG-STAR, MLST, NG-MAST and NG-MAST genogroups. The colour of each lineage matches that in the coloured strips in (a). The tree and typing information obtained by pyngoST from this dataset can be explored using Microreact (https://microreact.org/project/wYpBzCs9A6Uf7HEMA6zmmY-eurogasp2018-pyngost).

We also evaluated the distribution of pairwise differences among the core genome of isolates from the same lineage (ST, CCs and genogroups) independently of the phylogeny. The SNP distance in the core genome of all the isolates in the dataset ranged from 0 to 3,883. The distribution of pairwise SNPs among isolates from the top lineages of the five schemes clearly revealed the existence of genomic sublineages within the same ST, CC or genogroup (Figure 2b). At a short scale, 69.7% of isolates of the same NG-MAST STs, followed by 60.0% of isolates of the same NG-STAR ST and 43.3% isolates of the same MLST ST, differed in less than 100 SNPs between each other. NG-STAR CCs and NG-MAST genogroups, which group different STs, reduce these proportions to 30.6% and 58.0%, respectively. At a longer scale, 99.5% of NG-STAR STs (82.9% of CCs) are formed by isolates below 1,000 SNPs, followed by 96.8% of NG-MAST STs (94.2% of genogroups) and 90.5% MLST STs.

## CONCLUSIONS

Traditional molecular typing tools are still widely used for classifying gonococcal isolates into STs for epidemiological and surveillance studies. However, WGS is increasingly being used for these purposes as it provides the maximum discrimination and accuracy in resolution between isolates. From these genomes, we can directly extract the typing information, avoiding tedious amplification and sequencing reactions and contributing to the integration and comparability of results from epidemiological and surveillance studies that use WGS with those that use traditional molecular typing. pyngoST is a fast command-line Python tool for rapid and accurate multiple sequence typing of *N. gonorrhoeae* genome assemblies using any or all of the main typing schemes available: MLST, NG-MAST and NG-STAR. It can also output NG-STAR CCs for an easier representation of AMR lineages and calculate NG-MAST genogroups for reproducibility of results from previous studies. Genogroups are dependent on the dataset and change over time and thus, their use is not encouraged, in benefit of other schemes such as NG-STAR STs and CCs, which are comparable among studies. Finally, we show that NG-STAR is the scheme that best represents the genomic population structure of *N. gonorrhoeae*, highlighting the importance of antimicrobial use and AMR on the evolution of this sexually-transmitted pathogen.

## Author contributions

LSB, DG and MU designed the study. LSB, ASS and DG wrote the code and tested intermediate and final versions. LSB wrote the first draft of the manuscript and ASS, DG and MU read, commented on, and approved the final manuscript.

## Acknowledgements

We thank Simon R. Harris for the initial code on calculating NG-MAST genogroups used in the Euro-GASP 2013 genomic survey (8).

